# Aryl hydrocarbon receptor knockout accelerates PanIN formation and fibro-inflammation in a mutant *Kras*-driven pancreatic cancer model

**DOI:** 10.1101/2023.02.01.526625

**Authors:** Morgan T. Walcheck, Patrick B. Schwartz, Noah D. Carrillo, Kristina A. Matkowsky, Manabu Nukaya, Christopher A. Bradfield, Sean M. Ronnekleiv-Kelly

**Affiliations:** University of Wisconsin School of Medicine and Public Health, Department of Surgery, Division of Surgical Oncology, K4/747 CSC, 600 Highland Avenue, Madison, WI 53792; University of Wisconsin, McArdle Laboratory for Cancer Research, 1400 University Avenue, McArdle Research Building, Madison, WI, 53706; University of Wisconsin School of Medicine and Public Health, Department of Pathology and Laboratory Medicine, L5/183 CSC, 600 Highland Avenue, Madison, WI 53792; University of Wisconsin Carbone Cancer Center, Madison, WI 53705; William S. Middleton Memorial Veterans Hospital, Madison, WI 53705

**Keywords:** Pancreatic Cancer, Aryl hydrocarbon Receptor, Pancreatic intraepithelial neoplasia (PanIN), Fibro-inflammatory infiltrate, Single Cell RNAseq, Aurora (Cytek)

## Abstract

**Objectives:** The pathogenesis of pancreas cancer (PDAC) remains poorly understood, hindering efforts to develop a more effective therapy for PDAC. Recent discoveries show the aryl hydrocarbon receptor (AHR) plays a crucial role in the pathogenesis of several cancers, and can be targeted for therapeutic effect. However, its involvement in PDAC remains unclear. Therefore, we evaluated the role of AHR in the development of PDAC *in vivo*.

**Methods:** We created a global AHR-null, mutant *Kras*-driven PDAC mouse model (A^-/-^KC) and evaluated the changes in PDAC precursor lesion formation (Pan-IN 1, 2, and 3) and associated fibro-inflammation between KC and A^-/-^KC at 5 months of age. We then examined the changes in the immune microenvironment followed by single-cell RNA-sequencing analysis to evaluate concomitant transcriptomic changes.

**Results:** We found a significant increase in PanIN-1 lesion formation and PanIN-1 associated fibro-inflammatory infiltrate in A^-/-^KC vs KC mice. This was associated with significant changes in the adaptive immune system, particularly a decrease in the CD4+/CD8+ T-cell ratio, as well as a decrease in the T-regulatory/Th17 T-cell ratio suggesting unregulated inflammation.

**Conclusion:** These findings show the loss of AHR results in heightened *Kras*-induced PanIN formation, through modulation of immune cells within the pancreatic tumor microenvironment.

## INTRODUCTION

Pancreatic ductal adenocarcinoma (PDAC) is one of the most lethal cancers and is currently the fourth leading cause of cancer-related deaths in the United States.^1^ In 2022, 49,830 out of 62,210 (80%) patients diagnosed with PDAC died of their disease.^1^ These dismal outcomes are a consequence of a poor understanding of PDAC pathogenesis, which continues to hinder efforts to develop more effective prevention strategies as well as treatments for patients with PDAC. To develop such therapies, it is essential to elucidate key pathways involved in the development of PDAC that can then be exploited for preventive or therapeutic effect. Recent discoveries show the aryl hydrocarbon receptor (AHR) transcription factor plays a crucial role in the development of several cancers through mediation of immune cell infiltration and function and holds promise as a target in PDAC.^2–6^

The aryl hydrocarbon receptor (AHR) is a ligand-activated transcription factor that is a member of the basic helix-loop-helix (bHLH) Per-Arnt-Sim (PAS) superfamily.^7^ It was traditionally characterized as a xenobiotic receptor for sensing environmental pollutants, such as 2,3,7,8-tetrachlorodibenzo-p-dioxin and benzo(a)pyrene, but recently it has been shown to be involved in many endogenous cellular processes whose dysregulation leads to cancer development.^7^ Although the exact role the AHR plays in the development of cancer is unclear, it is suspected that AHR may mediate tumorigenesis through its role in inflammation and immune cell regulation ^2,3,8–10^. For instance, the AHR is known to influence immune cell development such as the differentiation of T-helper (Th) cells or T-regulatory (T-reg) cells.^11^ Furthermore, the AHR modulates resident immune cells in epithelial tissues and has an emerging role in immunosurveillance.^4^ While there is compelling data elucidating the role of AHR in several different types of cancer^2,6,8^, in PDAC, there is promising but sparse data suggesting a tumor suppressor function.^12,13^ For example, previous studies have shown improved survival in PDAC patients with high expression of AHR in cancer tissue compared to patients with low AHR expression.^12^ Furthermore, when studied in established PDAC cell lines, increased activation of AHR demonstrated inhibition of cancer cell growth while blocking AHR activation increased growth.^12^ These data suggest a potential tumor suppressive role of AHR in PDAC but this question has not been evaluated *in vivo* nor has the impact of the AHR on the PDAC tumor microenvironment (TME) been assessed.

Therefore, in the current study, we sought to understand how AHR modulates the developing TME to impact PDAC formation *in vivo*. We hypothesized that loss of AHR would cause accelerated pancreatic cancer precursor lesion formation (pancreatic intraepithelial neoplasia or PanIN) due to changes in the immune microenvironment. Using a well-established model of pancreas cancer precursor lesion development (*LSL-Kras^G12D/+^; Pdx1-Cre* or KC mice), we generated a global *AHR* knockout in the pancreas-lineage *Kras*-mutant mouse (A^-/-^KC). We evaluated A^-/-^KC mice at 5 months of age, a timepoint where early PanIN formation occurs and the TME begins to develop, which would enable us to determine how loss of AHR may modulate development and progression of the pancreatic cancer precursor (PanIN) lesions and the TME. We found knockout of AHR in the KC mice resulted in distinct changes in the TME, specifically we identified accelerated formation of PanIN-1 and an increase in the surrounding fibro-inflammatory infiltrate. To understand what changes the knockout of the AHR induced in this region we analyzed the changes in immune cell populations within the A^-/-^KC pancreas compared to the KC pancreas. We found a significant depletion of CD4+ T-regs and a decreased CD4+/CD8+ T-cell ratio in the A^-/-^KC mouse pancreas compared to the KC mouse pancreas, suggesting diminished inflammatory regulation as a consequence of AHR loss. We further evaluated the transcriptomic changes induced by loss of AHR through single cell RNA sequencing (scRNAseq) and found changes in gene expression within T-cell clusters indicating changes in T-cell function after AHR knockout. Overall, these findings uncover the integral role of AHR in *Kras*-induced PDAC pre-cursor lesion formation and mark it as a potential target in PDAC that can be exploited for preventive or therapeutic effect.

## METHODS

### Animals

All animal studies were conducted according to an approved protocol (M005959) by the University of Wisconsin School of Medicine and Public Health (UW SMPH) Institutional Animal Care and Use Committee (IACUC). Mice were housed in an Assessment and Accreditation of Laboratory Animal Care (AALAC) accredited selective pathogen-free facility (UW Medical Sciences Center) on corncob bedding with chow diet (Mouse diet 9F 5020; PMI Nutrition International), and water ad libitum. The Lox-Stop-Lox (LSL) *Kras^G12D^* (B6.129S4-*Kras* tm4Tyj/J #008179), and *Pdx1-Cre* (B6.FVB-Tg(*Pdx1-Cre*)6Tuv/J) mice were purchased from the Jackson Laboratory (Bar Harbor, ME). The global *AHR* knockout (*AHR^-/-^*) mice were a gift from the laboratory of Chris Bradfield. All mice were housed under identical conditions and are congenic on a C57BL/6J background (backcrossing > 15 generations). The *LSL-Kras^G12D^* and *Pdx1-Cre* mice were bred to develop *LSL-Kras^G12D^*; *Pdx1-Cre* (KC) mice. The KC mice were bred to the *AHR^-/-^* mice to develop the *AHR^+/--^; LSL-Kras^G12D^*; *Pdx1-Cre* (A^+/--^KC) and the AHR-null; *LSL-Kras^G12D^*; *Pdx1-Cre* (A^-/-^KC) mice. Genotyping for *Kras* and *Pdx1-Cre* was performed according to Jackson Laboratory’s protocols (Cre: Protocol #21298, *Kras^G12D^:* Protocol #29388, AHR protocol #20775) and activated Kras based first on a modified protocol published by Hingorani.^15–17^ The health and well-being of the mice were monitored closely by research and veterinary staff. Mice that showed signs of distress, such as significant weight loss (> 10% total body weight), poor feeding, poor mobility, hunched back, or significant hair loss, were immediately euthanized. Mice were euthanized through CO^2^ asphyxiation.

### Histology

Pancreas specimens were collected and formalin-fixed and paraffin embedded (FFPE) by standard methods. Any external tumors (facial tumors and anal tumors), including surrounding normal tissue, were also removed for pathologic analysis. Pancreas, liver, spleen, kidney and external samples were sectioned at 4 um thickness and 200 um intervals for a total of 10 slides. Sectioning and hematoxylin and eosin (H&E) staining was performed by the University of Wisconsin Experimental Animal Pathology Lab (EAPL) core facility.

### Pancreatic intraepithelial neoplasia (PanIN) and cancer evaluation

The mice in this study were evaluated at 5 of age. At the time of euthanization, mice underwent cervical dislocation followed by midline laparotomy. A board-certified surgical pathologist with subspecialty training in gastrointestinal pathology (KM) who was blinded to the mouse genotype and sex evaluated the pancreatic histology sections. All resected organs were evaluated for abnormal pathology such as cancer development. Each pancreas was evaluated for extent of involvement of fibrosis / immune cell infiltrate (fibro-inflammatory infiltrate) and the extent / grade of pancreatic intraepithelial neoplasia (PanIN 1-3). The presence of PDAC was recorded along with any other noted pathology. Comparisons of tumor development between groups was accomplished using the Fisher’s Exact test. Data was considered significant with a p-value <0.05. All statistics were done using GraphPad.

### Single cell dissociation of the pancreas

Mice were sacrificed using cervical dislocation and the pancreas was removed and placed in ice cold PBS in under 2 minutes. The pancreas was then minced for 60 seconds, washed in 10ml PBS and spun at 300g and 4°C for 5 minutes. Following a second PBS wash and centrifugation, the pancreas was placed in a 15 mL conical tube in a digestion buffer containing 1 mg/mL collagenase P dissolved in HBSS. The conical tube was submerged horizontally in water at 36°C, and the pancreas was digested for 15-30 minutes, manually agitating every 5 minutes. After approximately 15 minutes of digestion, the reaction was quenched with 10 mL ice cold R10 media (RPMI and 10% FBS). The pancreas was then spun down and washed with R10 media, and subjected to a filter gradient (500, 100 and 40 um filters). The resultant single cell suspension was then washed and resuspended in a flow buffer (PBS and 2% FBS).

### Flow cytometry

Single-cell suspensions were washed and resuspended in flow buffer. To determine cell viability, cells were first stained with LIVE/DEAD Fixable Blue Dead Cell Stain Kit (ThermoFisher Scientific) according to manufacturer instructions. The cells were washed and then incubated for 30 mins at 4°C in the antibody cocktail for surface staining. The list of flow antibodies is provided in supplemental table 1. Finally, the cells were washed and then FoxP3 intracellular staining was performed using the FoxP3/Transcription Factor Staining Buffer Set (eBioscience) according to manufacturer instructions. Flow cytometry of samples was performed with an Aurora spectral flow cytometer (Cytek). Flow cytometry data was analyzed using the gating strategy in Supplemental Figure 1 in FCS express 7 (DeNovo software). Comparisons of immune cell changes between each genotype was done using the Fisher’s Exact test. Data was considered significant with a p-value < 0.05. All statistics were done using GraphPad.

### Single-cell RNAseq

After the pancreas was dissociated into single cells, the cells were stained with DAPI (1:1000) and then live/dead sorted into single cell populations into PBS + 2%FBS. These were transferred to the University of Wisconsin Gene Expression Center for Library Preparation. In brief, libraries were constructed according to the Chromium NextGEM Single Cell 3’ Reagent Kit v3.1 User Guide, Rev.D (10x Genomics, Pleasanton, CA). Single cell suspension cell concentration and viability were quantified on the Countess II (Thermo Fisher Scientific, Waltham, MA) using 0.4% Trypan Blue (Invitrogen, Carlsbad, CA). Next, 10 x10^3^ cells were mixed with 10X Genomics Chromium single-cell RNA master mix, followed by loading onto a 10X Chromium chip according to the manufacturer’s protocol to obtain single-cell cDNA. Libraries were sequenced on a NovaSeq6000 (Illumina, San Diego, CA).

Single cell RNAseq data were analyzed by the UW Bioinformatics Resource Center. Data were demultiplexed using the Cell Ranger Single Cell Software Suite, mkfastq command wrapped around Illumina’s bcl2fastq. The MiSeq balancing run was quality controlled using calculations based on UMI-tools.^18^ Samples libraries were balanced for the number of estimated reads per cell and run on an Illumina NovaSeq system. Cell Ranger software was then used to perform demultiplexing, alignment, filtering, barcode counting, UMI counting, and gene expression estimation for each sample according to the 10x Genomics documentation (https://support.10xgenomics.com/single-cell-geneexpression/software/pipelines/latest/what-iscell-ranger). The gene expression estimates from each sample were then aggregated using Cellranger (cellranger aggr) to compare experimental groups with normalized sequencing-depth and expression data.

Gene expression data was then processed using the Seurat version 3 pipeline for data integration.^19^ Data was loaded into Seurat. Low quality (<200 features) and doublets were removed. Cells expressing more than 1% hemoglobin gene content were removed. The KC and A^-/-^KC data was merged and integrated using the SCTransform wrapper with 3000 features.^20^ SCTransform allowed for normalization and better batch correction based on a regularized negative binomial regression. The regression allowed for correction of the cell cycle effect and mitochondrial gene content. A principal component analysis (PCA) was performed with the number of dimensions (principal components) selected based on an elbow plot. Data was visualized using the Uniform Manifold Approximation and Projection for Dimension Reduction (UMAP).^21^ Cell identities were called based on the expression of known markers for each cell type. Differential expression between KC and A^-/-^KC was performed between clusters in Seurat using MAST test based on a log fold change threshold of 0.25 and an adjusted p value of p < 0.05 and visualized using heatmaps generated with the DoHeatmap function within Seurat. The MAST test within Seurat is based on the MAST package, which is specifically designed to handle differential expression in scRNA-seq data and utilizes a hurdle model.^22^ Differences between the proportion of cells was compared using a custom code (https://github.com/rpolicastro/scProportionTest/) based on permutations test which compares the proportions of cells in each cluster between two scRNA-seq samples. Data was presented as a p value with an associated confidence interval as a point range plot. Differences with a false discovery rate q < 0.05 and absolute log2 fold difference of > 0.58 were considered statistically significant. Gene Set Enrichment Analysis (GSEA) was performed by converting all mouse gene symbols to their human homologs with biomaRt.^23,24^ The fgsea package^25^ was then used to identify enriched pathways within the ImmuneSigDB subset of C7 MSigDB collection^26,27^ with the following settings: minSize = 15, eps = 0.0, maxSize = 500, and an adjusted p-value of < 0.05 for CD4+ T-cells.

## RESULTS

### Extent of PanIN and associated fibro-inflammatory infiltrate is significantly increased in A^-/-^KC mice at 5 months of age

We evaluated the pancreas histology of KC, A^+/--^KC and A^-/-^KC at 5 months of age. We found a statistically significant increase in PanIN-1 and associated fibro-inflammatory infiltrate in A^-/-^KC compared to KC mice (figure 1A, C). This finding of accelerated formation of PanIN-1 and associated fibro-inflammation is relevant for tumor promotion due to an increase in the amount of pancreas ‘at risk’ for progression to PanIN-2, 3 and PDAC. Although not statistically significant, we did identify high grade PanIN (PanIN-3) and PDAC in the A^-/-^KC mice (n = 1 each) at age 5 months, an observation that has not been reported or identified in KC mice which mostly exhibit normal pancreas at this time.^15^ These results indicate that knockout of AHR in KC mice accelerates PanIN-1 / fibro-inflammatory infiltrate formation.

**Figure 1:**
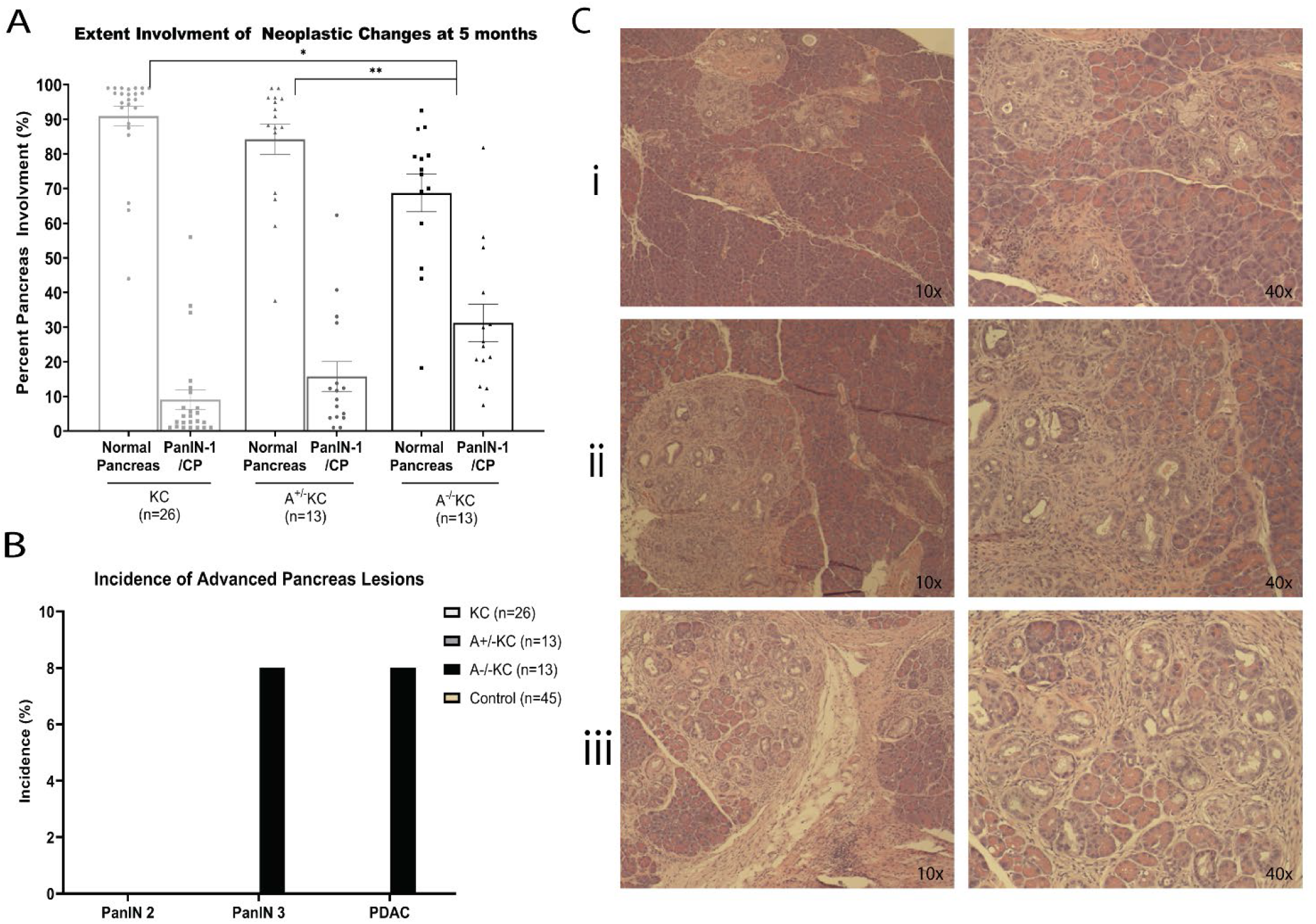
Extent of PanIN and associated fibro-inflammatory infiltrate is significantly increased in A^-/-^KC mice at 5 months of age. At 5 months, there is a significant increase in extent of pancreas undergoing neoplastic changes in the A^-/-^KC mice compared to KC and A^+/--^ KC. At 5 months there is a significant increase in the extent of pancreas involvement with 9% and 15% of KC and A^+/--^KC compared to 31% of A^-/-^KC pancreas afflicted (KC vs A^-/-^KC P-value = 0.0017, A^+/--^KC vs A^-/-^KC P-value =0.02). There is no advanced pathology seen in KC and A^+/--^ KC mice (B), but there are occurrences of high-grade PanIN-3 and PDAC in the A^-/-^KC group (B). This is not statistically significant, although an instance of PanIN-3 nor PDAC has not been reported in the KC model at this age. Representive images of the percent pancreas with pathology in KC (Ai), A^+/--^KC (Aii) and A^-/-^KC (Aiii) at 10x and 40x respectivly. KC and A^+/--^KC mice have on average 9% and 15% extent pancreas involvment respectivly (Ci, Cii). A^-/-^KC mice had an average of 31% extent involvment (Ciii).

### Significant increase in CD45+ immune cell infiltration into the pancreas after AHR knockout

To understand what could be driving the increase in PanIN-1 lesions and associated fibro-inflammatory infiltrate (i.e. tumor promotion) after *AHR* knockout, we first evaluated changes in populations of CD45+ infiltrating immune cells within the pancreas between C57BL/6J mice, A^-/-^, KC and A^-/-^KC mice. The AHR is expressed within most immune cell populations with highest expression in Th17 cells, T-regulatory cells, macrophages, and DCs.^28^ The AHR has also been shown to influence function of immune cells and interaction with epithelial cells through several different mechanisms depending on the organ system, so the global knockout of AHR could induce a variety of changes in the infiltrating immune cells. Concordantly, we found an increase in CD45+ immune cells in the A-/- mice, compared to C57BL/6J mice (Figure 1). The level of CD45+ infiltration in the A-/- pancreas was equal to that in the KC and A-/-KC groups despite the lack of a *Kras*-mutation and disease development in the A-/- pancreas. Furthermore, although an increase in CD45+ immune cells was identified in the A-/- pancreas versus C57BL/6J mice, pancreata between these two groups were histologically indistinct suggesting the knockout of *AHR* elicits changes in the immune infiltration into the pancreas irrespective of an underlying pathogenic process.

### Increase in PanIN-1 lesions after AHR knockout is not due to innate immune cells

An immunosuppressive microenvironment is highly characteristic in PDAC development.^29,30^ Infiltrating immune-suppressor cells, such as myeloid derived suppressor cells (MDSCs), neutrophils, and macrophages, are found around the primitive PanIN-1 lesions and directly promote acinar cell dedifferentiation during the earliest stages of pancreatic pre-cancer formation. ^31–35,36^ We found no change in MDSCs and neutrophils between A^-/-^KC and KC mouse pancreas suggesting these groups are not influencing the increase in PanIN lesions (Supplemental Figure 2). Interestingly, we found that the knockout of AHR in both in wild-type and *Kras*-mutant pancreas resulted in a significant decrease in the macrophage population (Figure 3). Macrophages have been shown to increase PanIN formation and progression so an increase in macrophages is a common finding in PDAC and typically associated with advanced PDAC.^36^ Finding a decrease in the macrophage population, along with no changes in MDSCs or neutrophils, suggests that the increase in PanIN-1 formation after AHR knockout is unlikely attributed to changes in the immunosuppressive immune types typically implicated in influencing PDAC development.

**Figure 2:**
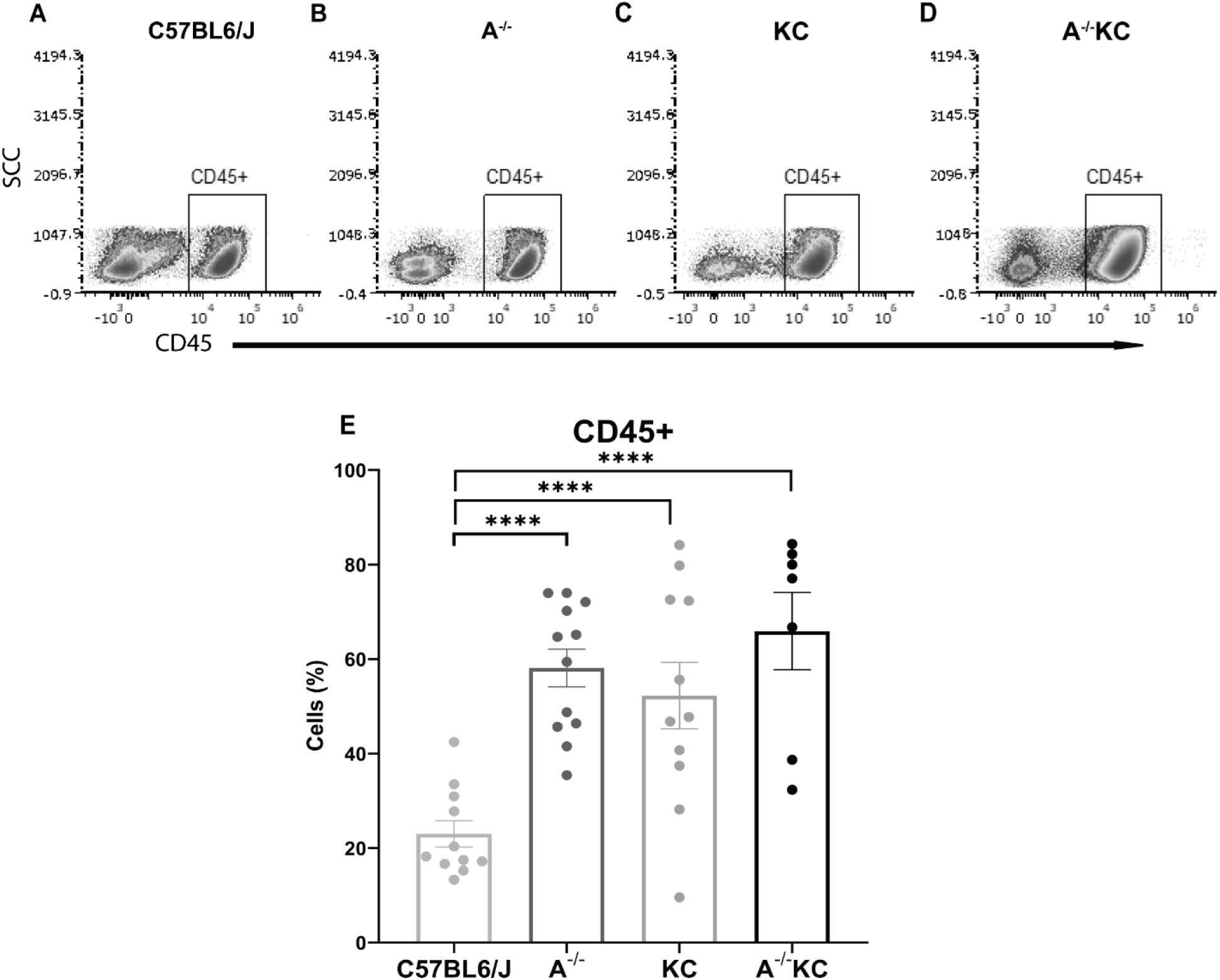
Significant increase in CD45+ cells in A^-/-^, KC and A^-/-^KC pancreas compared to C57BL/6J. There was a statistically significant increase in CD45+ cells, a broad immune cell marker, in A^-/-^, KC and A^-/-^KC pancreas compared to C57BL/6J (58.14%, 52.19%, 65.94% vs 23.04% respectively) (all P<0.0001). There were no statistically significant differences in CD45+ cells between A^-/-^, KC and A^-/-^KC pancreas.

**Figure 3:**
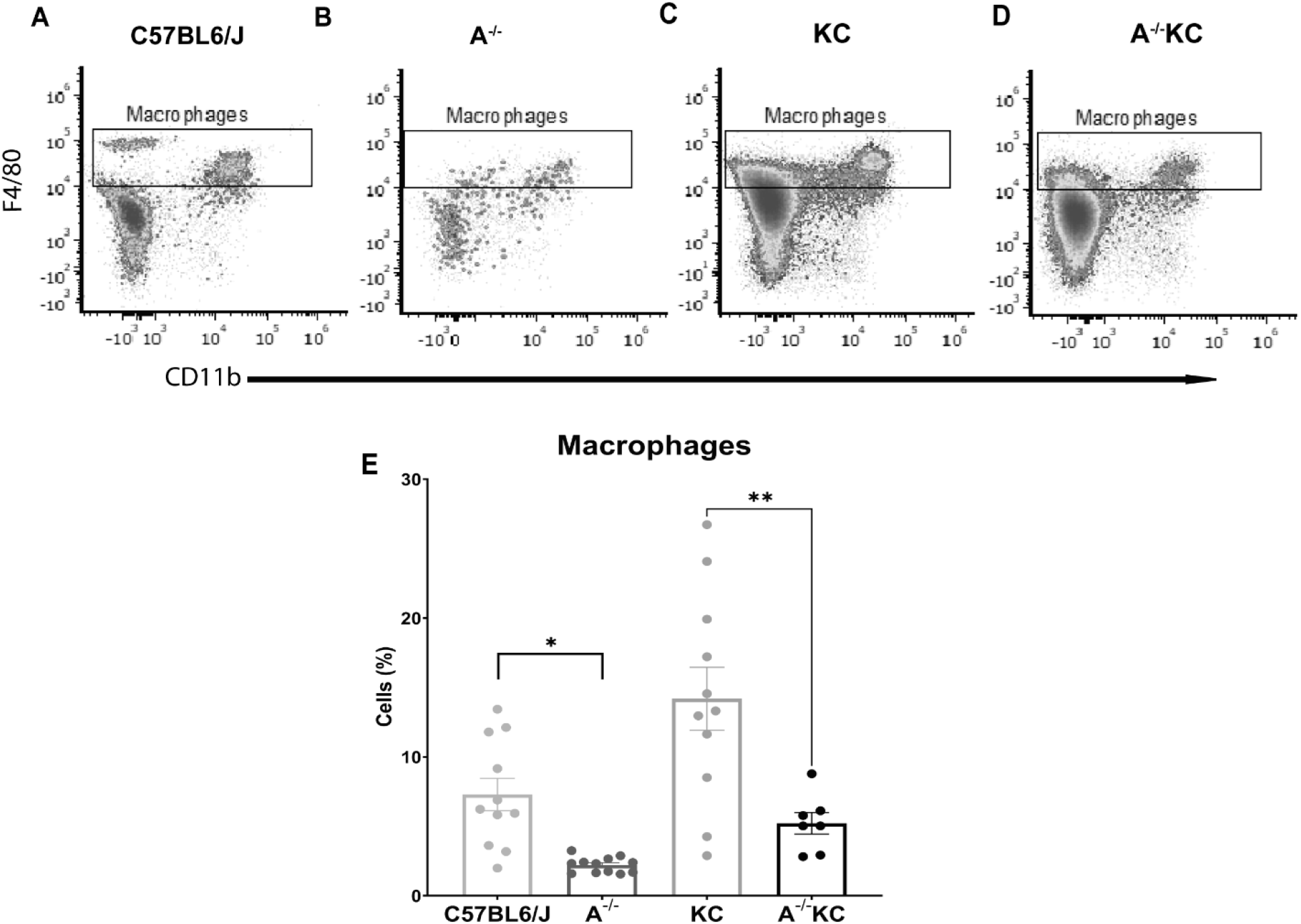
Knockout of AHR results in a decrease in macrophages. The knockout of AHR in healthy pancreas resulted in a decrease in macrophages in A^-/-^ compared to C57BL6/J (2.210% vs 7.93%, p-value = 0.0014) as well as in A^-/-^KC pancreas compared to KC pancreas (5.216% vs 14.19%, p-value =0.0027).

In addition to increases in immunosuppressive immune types, there is usually a notable lack of dendritic cells (DCs) and natural killer (NK) cells found in the PDAC microenvironment.^37^ The number of DC and NK cells are lowest in the pancreas of patients with a tumor, while exogenously increasing the number of DC or NK cells has been shown to reduce PDAC tumor and PanIN size.^38^ However, in this study, we found no changes in DCs (Supplemental Figure 2) and a significant increase in NKs after AHR knockout in KC mice (Figure 4) suggesting the increase in PanIN-1 formation should be attributed to another immune group.

**Figure 4:**
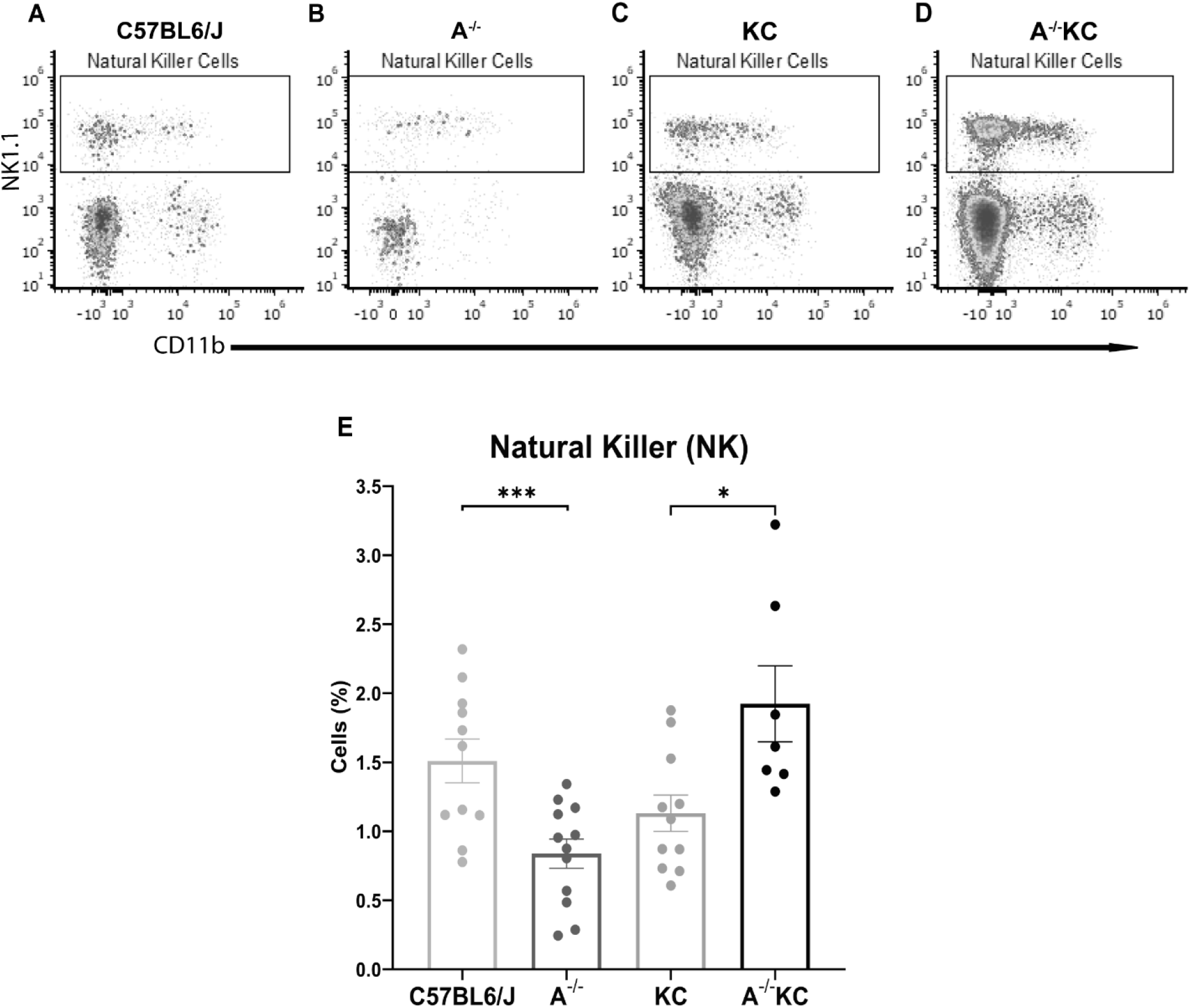
Significant increase in natural killer (NK) cells in A^-/-^KC pancreas. There was a significant decrease in NK cells in A^-/-^ pancreas (B) compared to C57BL6/J (A) (1.5% vs 0.75%, P-value <0.0001), but an increase in NK cells in the A^-/-^KC (D) pancreas compared to KC pancreas (C) (1.1% vs 1.8%, P-value <0.01).

### Changes in adaptive immune cells after AHR knockout increase PanIN-1 lesion formation

Due to the established role of AHR in modulating the development of adaptive immune cells particularly T-cells, we evaluated for changes in B-cells, CD8+ T-cells, CD4+ T-regulatory (Treg) cells and CD4+ T-helper 17 (Th17) cells. We analyzed CD4+ and CD8+ T-cells using the CD4+/CD8+ ratio. CD4+ T-cells help coordinate immune function by stimulating and/or regulating other immune cell response. Cytotoxic CD8+ T-cells kill infected cells, cancerous cells, or damaged cells, and consequently induce inflammation. A diminished CD4+/CD8+ ratio indicates decreased regulatory cells compared to cytotoxic T-cells, which corresponds to heightened inflammation. In the A^-/-^KC pancreas, we identified a significant decrease in the CD4+ T-cell population resulting in a decrease in CD4+/CD8+ ratio compared to KC mouse pancreas (0.26 → 0.54) (Figure 6I, J). This low ratio is typically seen in the setting of pancreatic tissue inflammation.^39^ The decrease in CD4+ T-cells was primarily due to a significant decrease in FoxP3+ T-regs. CD4+ Th17 cells were unchanged (Figure 6K, L). Of note, the A^-/-^ mouse demonstrated a significant increase in CD8+ T-cells compared to C57BL/6J mice, but there was a concomitant substantial increase in CD4+ T-cells, thus balancing out the CD4+/CD8+ ratio. Together, this data indicates that in the setting of an ongoing insult such as the mutant-*Kras* driven PanIN / fibro-inflammatory infiltrate formation, loss of AHR results in a significant decrease in anti-inflammatory regulation (decreased CD4+ T-regs), which may exacerbate the phenotype.

**Figure 5:**
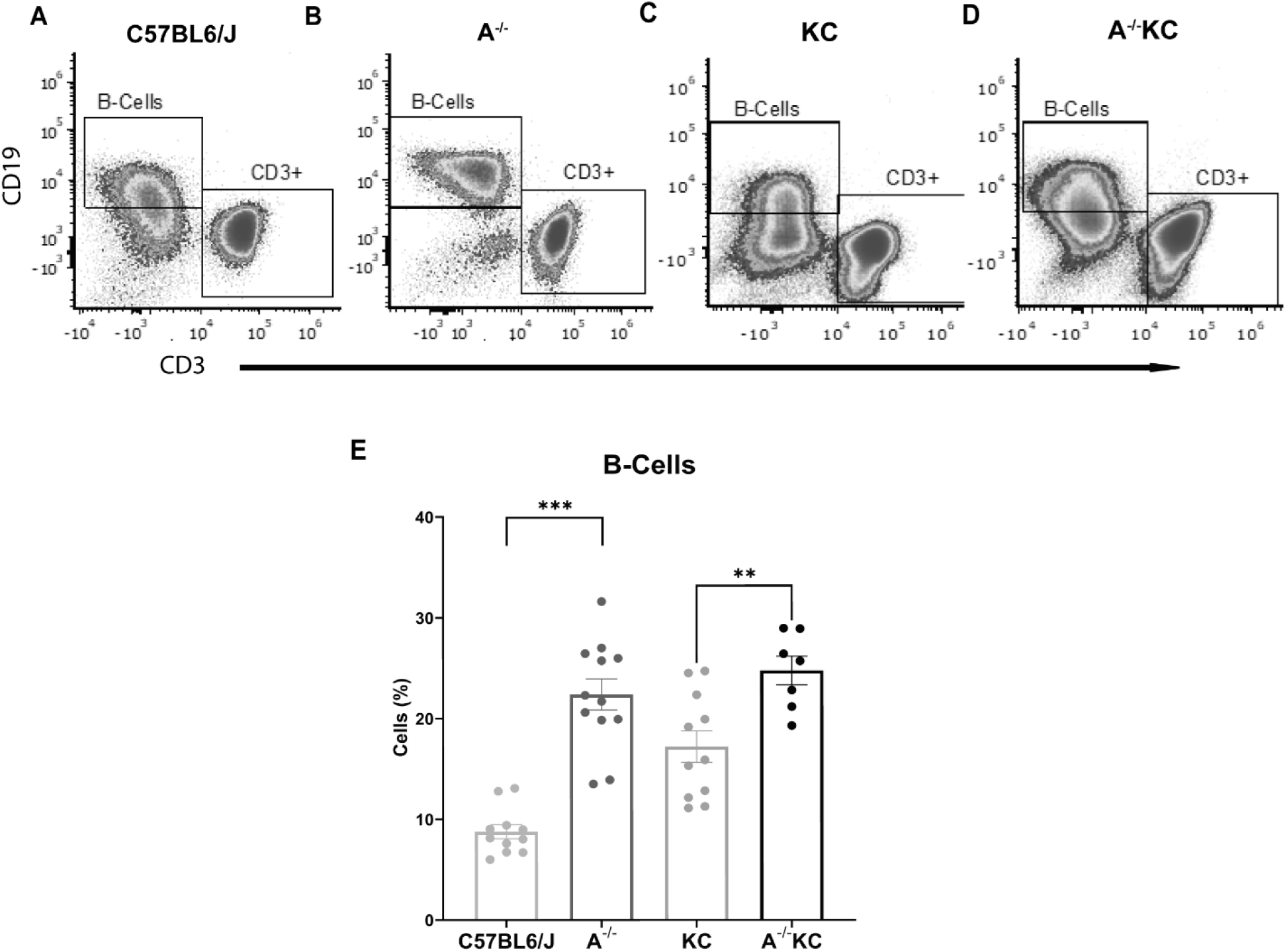
AHR knockout results in a significant increase in pancreatic B-cells. There is an increase in infiltrating B-cells in A^-/-^ pancreas compared to C57BL/6J mice (22.38% vs 8.77%, p < 0.0001). B-cell infiltration also increased in A^-/-^KC mice vs KC mice (24.21% vs 18.07%, p = 0.0416).

**Figure 6:**
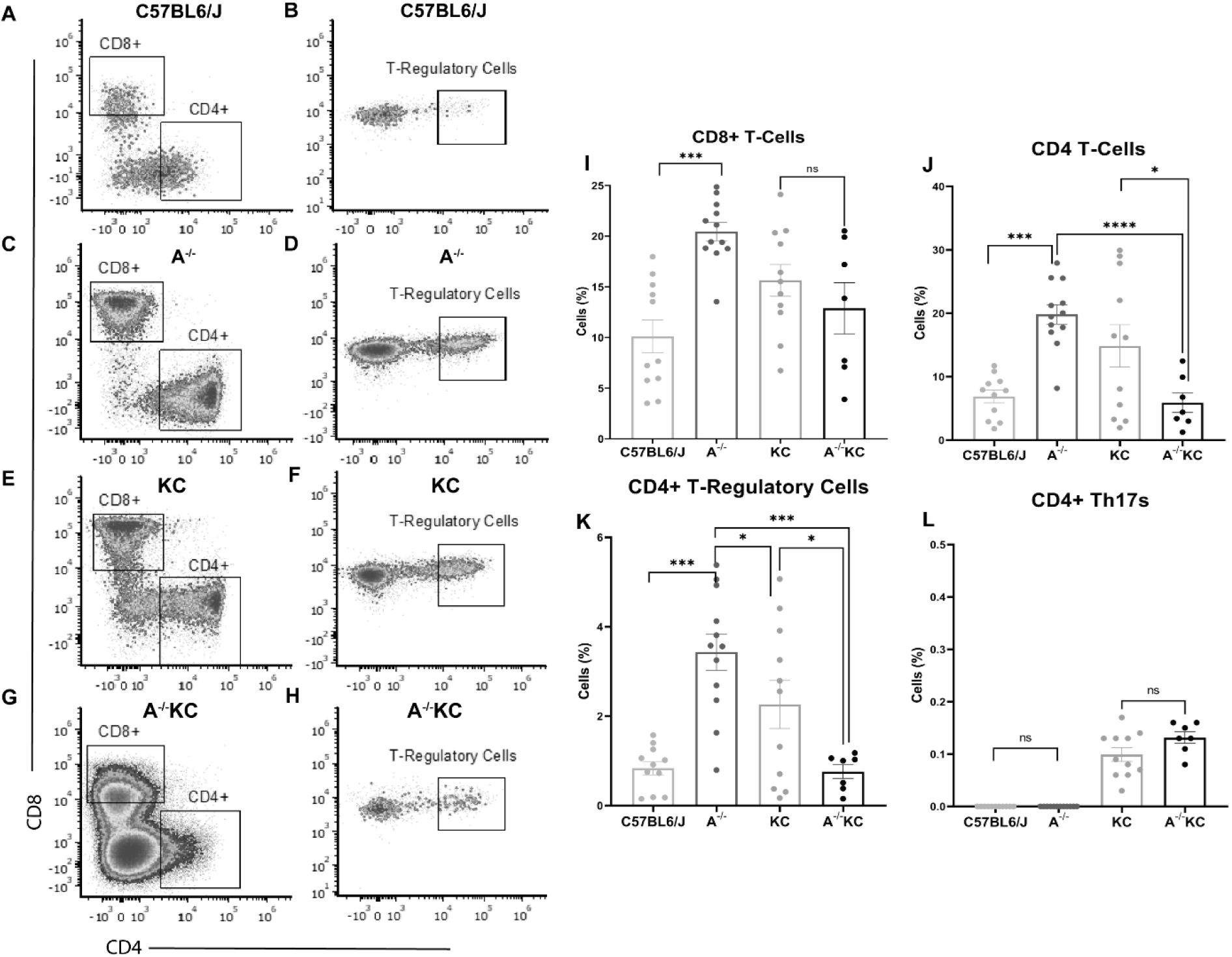
Significant decrease in immunosuppressive T-cell groups in A^-/-^KC pancreas. There was a significant increase in CD4+ T-cells in A^-/-^ pancreas compared to C57BL6/J (19.82% vs 6.889%, P-value < 0.0001), but a decrease in CD4+ cells in A^-/-^KC pancreas compared to KC pancreas (5.901% vs 14.86%, p-value = 0.0291) (A). There was a significant increase in CD8+ T-cells in A^-/-^ pancreas vs C57BL6/J (20.45% vs 10.10%, p-value < 0.0001), but there were no differences between KC and A^-/-^KC pancreas (B). There was a significant increase in T-regs in A^-/-^ pancreas compared to C57BL6/J (3.431% vs 0.8390%, P-value < 0.0001) and a significant decrease in A^-/-^KC pancreas compared to KC pancreas (0.7606% vs 2.261%, p-value = 0.0202) (C). There were no differences in Th17 cells in A^-/-^ compared to C57BL6/J or in A^-/-^KC compared to KC pancreas (D).

In addition to significant increases in T-cell populations, we found a significant increase in B-cells after AHR knockout in both WT and KC mice (figure 5). The role AHR has in B-cell development, proliferation and function as well as the role B-cells play in cancer are new and developing areas of study.^40,41^ In PDAC, It has been shown that the progression of PanIN lesions is associated with an increased amount of B-cell infiltration within the pancreas of both patients and mice with PDAC.^42^ Heightened B-cell infiltration has also been linked to worsened prognosis and increased fibrosis within the pancreas.^41^ The increase in B-cells after AHR knockout suggests B-cells may also be directly involved in the increase in PanIN-1 formation.

### An Increase in immune cell populations and a decrease in fibroblasts found in the A^-/-^KC pancreas with single-cell RNA sequencing

To further understand cellular changes that occur with loss of AHR, particularly to gain insight into what is changes occur transcriptionally in the infiltrating immune cells, we performed single cell RNA sequencing comparing the A^-/-^KC pancreas to the KC pancreas. Other cell populations such as fibroblasts can contribute to PanIN progression and PDAC development, so we first looked for any changes within this population in the A^-/-^KC pancreas. Overall, there did not appear to be evidence supporting increased fibroblast populations that could drive accelerated fibrosis / PanIN formation in the A^-/-^KC pancreas (Supplemental Figure 3A, B). In fact, there appeared to be a decrease in these cell populations with a concomitant increase in immune cell populations including B-cells and NK cells (Supplemental Figure 1A, B), supporting the flow cytometry analysis. This also supported our hypothesis that changes in the immune system drive the increase in PanIN-1 formation after AHR knockout.

Next, given the significant decrease in CD4+ T-cells identified through flow cytometry, we analyzed the differential gene expression of the CD4+ T-cell cluster and performed a gene set enrichment analysis (GSEA) (Figure 7). GSEA revealed the CD4+ T-cell population had a significant upregulation of genes found in Treg vs Tconventional Thymus dataset suggesting Treg gene expression is downregulated in A^-/-^KC pancreas compared to KC pancreas (Figure 7B, C). The downregulated genes in the leading-edge signature (Odc1, Pim1, S100a4, Cxcr6, Dusp5, S100a6, Bhlhe40, Ccr2, Tmem176B, Fgl2, Dennd4a, Fosl2, Ifngr1, S100a10) and CD4+ differential gene expression heatmap further support this hypothesis (Figure 7D). These genes are often implicated in regulating the differentiation, function, cytokine production and survival of CD4+ T-cells.^43–47^ For example, Ornithine decarboxylase (ODC1) is the rate-limiting enzyme involved in the metabolism of polyamines and has been shown to be essential in regulating CD4+ T-cell differentiation.^43^ Overall, this suggests there are distinct changes in CD4+ T-cell differentiation after AHR knockout. This finding further corroborates the decrease in CD4+ T-cells and CD4+ T-cell subsets found using flow cytometry in this study.

**Figure 7:**
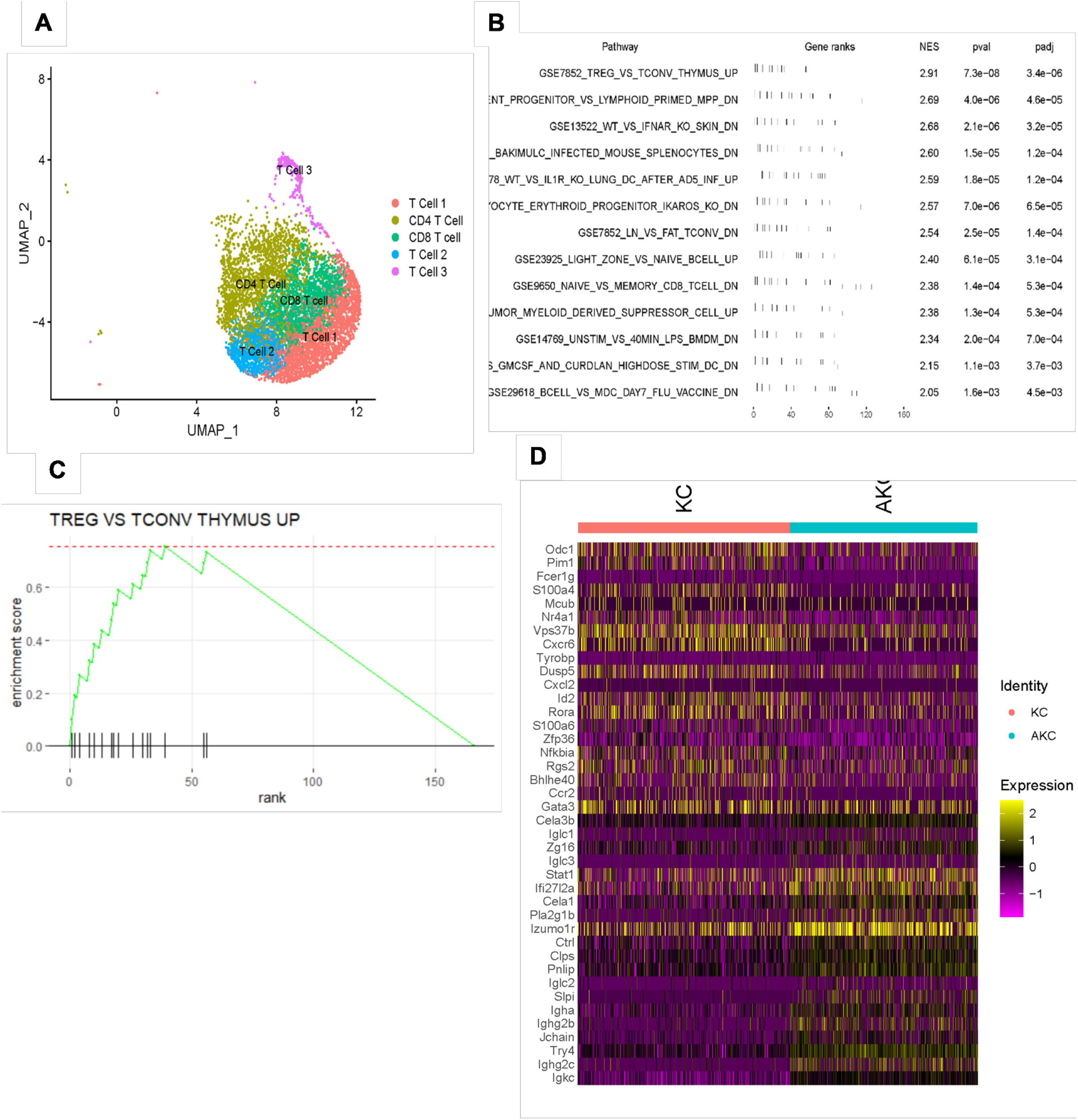
(A) The Uniform Manifold Approximation and Projection (UMAP) of the 5 T cell clusters identified during the scRNAseq analysis of KC and AKC pancreas. (B) Gene Set Enrichment Analysis (GSEA) pathway analysis on the CD4+ T-cell cluster demonstrating the pathways significantly different within the CD4+ T-cell cluster. (C) Enrichment score plot for the “TREG VS TCONV THYMUS UP” GSEA pathway (D) The CD4 T-cell cluster heat map showing the differential gene expression in the group that has a significant change in the GSEA.

## DISCUSSION

This study uncovered the integral role of AHR in modulating the development of pre-cancerous lesion formation of PDAC through its regulation of infiltrating immune cell populations in the fibro-inflammatory region. To evaluate the function of AHR in the development of PDAC we knocked out AHR from the KC PDAC pre-cursor mouse model and found a significant increase in PanIN-1 formation and associated fibro-inflammatory infiltrate at 5 months of age, correlating with the immune modulating role. This increase in pre-cancer lesion formation at 5 months suggests a higher likelihood of PDAC development given the greater amount of pancreas ‘at risk’ for progression. In support, Hingorani *et al* demonstrated that with increasing age, the extent of PanIN-1 replacement of pancreatic acinar cells in KC mice coincided with higher-grade PanIN-lesions. Along with the increase in PanIN-1 formation, we identified a significant increase in the associated fibro-inflammatory infiltrate with global AHR knockout leading us to hypothesize there is a potential immunologic influence causing accelerated PanIN lesions in this model.

To evaluate if there were any changes in infiltrating immune cell populations after AHR knockout, we utilized spectral flow cytometry to assess changes in MDSCs, neutrophils, macrophages, dendritic cells, natural killer cells, B-cells and T-cells. There has been mounting evidence that infiltrating immune cells found adjacent to PanIN lesions contribute significantly to lesion progression. One of the earliest immune responses documented after formation of early PanIN lesions is an infiltration of myeloid cells (MDSCs, neutrophils and macrophages) into PanINs that accumulate during PanIN development and PDAC formation.^31,32^ Myeloid cells directly promote acinar to ductal metaplasia (ADM) and progression of early PanIN lesions.^36^ For example, macrophages can secrete IL-6 and IL-10 which activate STAT3 resulting in increased PanIN progression through direct impacts on cell proliferation or through mediation of infiltrating cytotoxic immune cells.^36,48^ Moreover, MDSCs have been implicated as one of the main immune cell components contributing to PanIN progression through its promotion of an immunosuppressive environment. Interestingly, despite seeing an increase in PanIN-1 formation, we found no differences in neutrophils or MDSCs along with a significant decrease in the macrophage population in the A^-/-^KC mice. This suggests that the increase in PanIN-1 formation after AHR knockout is most likely not due to changes in immunosuppressive myeloid immune cells typically implicated in exacerbating PDAC development. In addition to typically finding increases in immunosuppressive immune types, there is usually a notable lack of dendritic cells (DCs) and natural killer (NK) cells found in the PDAC microenvironment. Typically, the number and functionality of DCs and NK cells decreases as PDAC progresses, but the reduction of NK cells begins with the formation of early PanIN lesions.^37,38^ Although we have an increase in PanIN-1 lesions, we found no changes in DCs and a significant increase in NK cells with loss of AHR suggesting the increase in pre-cursor lesions is more likely attributed to changes seen in the adaptive immune groups.

The presence of B-cells has been documented starting from early PanIN lesions through invasive cancer formation, and this population is thought to promote the progression of PanIN to cancer.^42^ Concordantly, existing data reveals that B-cells are increased around PanIN lesions of mouse model and human lesions.^42^ Recent studies have found B-cells support tumor growth through secretion of IL-35 which drives exclusion of CD8+ T cells (cells primarily responsible for immunosurveillance), increased fibrosis around PanIN lesions and tumor cell proliferation.^30^ This is further supported by results showing the depletion of B cells results in decreased PanIN formation in KC mice.^49^ After AHR knockout in the KC mouse model, we found a significant increase in B-cell infiltration which could further explain the increase in PanIN-1 formation seen in the A^-/-^KC mice. Unfortunately, there are limited studies depicting the specific role of B-cells in PDAC, and a lack of data demonstrating AHR impact on B-cell infiltration and function, but our study lends further evidence to the importance of B-cells in the progression of PDAC. Thus, these are currently developing areas of research.

In addition to innate immune cells and B-cells, the stroma of PDAC has a significant number of CD3+ T-lymphocytes that contribute to PanIN formation and progression through a variety of mechanisms.^50^ One subset of T-cells that has been shown to directly promote ADM and PanIN formation are CD4+ Th17 cells. Recent studies have shown infiltrating Th17 cells secrete the pro-inflammatory cytokine IL-17A inducing epithelial cell proliferation through their expression of the IL17A receptor thus promoting progression of ADM and formation of PanIN lesions.^51^ We found the knockout of AHR does not result in an increase in Th17 cells in the pancreas. Another T-cell subset implicated in promoting PanIN formation and progression are CD4+ T-regulatory cells (T-regs). Using mouse models, including the KC model, researchers have shown that immunosuppressive cells such as CD4+ FOXP3+ regulatory T cells (T-regs) accumulate both in PanIN and PDAC.^52^ A recent study found that T-regs facilitate the expansion of cancer associated fibroblasts (CAFs), which resulted in an increase in fibrosis around the PanIN lesions and PDAC.^52^ Conversely, the depletion of T-regs resulted in a decrease in fibrosis and an increase in PanIN formation.^52^ Despite the T-reg depletion, the immunosuppressive microenvironment was restored due to a consequent increase in myeloid suppressor cell infiltration around the PanIN lesions, which caused the PanIN progression.^52^ We found that AHR knockout resulted in a significant decrease in T-regs within the pancreas. Given the results from the previous study, the decrease in T-regs could further explain the increase in PanIN-1 formation. One key difference though, is their study showed an increase in other immunosuppressive cell types (MDSCs) while our study did not. This suggests that the increase in B-cells coupled with the decrease in T-regs may be responsible for the increased PanIN-1 formation.

Overall, the results of this study suggest the increase in PanIN-1 formation and surrounding inflammation after AHR knockout is mostly like due to changes within the adaptive immune cells versus a total increase in pro-inflammatory populations (or a decrease in other immunosuppressive cells such as MDSCs) typically seen in the pathology of PanIN development. Furthermore, our results suggest the increase in PanIN formation is primarily due to the changes within the infiltrating immune cells as opposed to other cell types in the PDAC microenvironment known to influence the formation and progression of PDAC. The PDAC microenvironment consists of immune cells as well as pancreatic stellate cells (PSCs) and cancer associated fibroblasts (CAFs) which are responsible for the deposition of the extensive extra cellular matrix (ECM) characteristic of PDAC.^53–55^ Each of these components have been shown to contribute to PanIN or PDAC formation or progression.^56–59^ PSCs are primarily responsible for the deposition of collagen 1 which has been shown to increase cancer cell proliferation, migration and resistance to apoptosis. CAFs have been shown to further drive the development of desmoplasia around the PanIN lesions through production of collagen types I and III, fibronectin, proteoglycans, and glycosaminoglycans. The increase in desmoplasia promotes tumorigenic phenotypes such as impaired drug penetration, poor vascularization, and reduced immune cell infiltration.^57,60,61^ Our results not only show a significant increase in immune cells, but also show a lack of changes within the other pancreatic parenchymal cell types. The results of the scRNAseq analysis corroborate the flow cytometry data while demonstrating a relative decrease in fibroblasts within the A^-/-^KC pancreas. Furthermore, the scRNAseq data identified Odc1 as one of the most significantly downregulated genes in A^-/-^KC according to the leading-edge signature (Figure 6). Odc1 regulates polyamine metabolism and is required for proper CD4+ T-cell differentiation into functional subsets.^43^ The knockout of Odc1 has been shown to disrupt CD4+ T-cell differentiation into functional subsets.^43^ The downregulation of Odc1 in A-/-KC mice indicates a decrease in CD4+ T-cell differentiation further corroborating the decrease in CD4+ T-cells and CD4+ T-cell subsets found in this study.

To our knowledge, this is the first study to depict a comprehensive analysis of AHR-dependent immune cell changes in pancreas. Our study showed the AHR plays an integral part in the heightened development of PDAC pre-cancer lesions, and correlates to changes in infiltrating immune cell populations. This phenotype is characterized by a significant increase in B-cells and a decrease in anti-inflammatory CD4+ T-cells, particularly FoxP3+ T-regulatory cells. The results of this study suggest the AHR plays a role in the early stages of PDAC development and in immune cell infiltration within the pancreas. Further work will investigate the effects of cell specific knockout of AHR, as well as AHR activation studies *in vivo*. Ultimately, given the ability of the AHR to control immune cell infiltration, the AHR holds potential as a possible target for anti-cancer immune modulation within the pancreas, particularly for patients who may be at high risk of developing PDAC (i.e. possess precursor lesions or strong family history).

## Acknowledgements

The authors thank the Experimental Animal Pathology Lab (EAPL) supported by the UWCCC (P30 CA014520) for use of its facilities and services.

