## Supplemental Figures for "Aryl hydrocarbon receptor knockout accelerates PanIN formation and fibro-inflammation in a mutant *Kras*-driven pancreatic cancer model"

**Supplemental Table 1:** Details about antibodies and fluorochromes used in flow cytometry.

| Fluorochrome | Target | Concentration (mg/ml) | Optimized volume to use (ul per 1 million per 100ul) |
|---|---|---|---|
| Live/Dead Blue | | | 1 |
| BUV737 | CD11c | 0.2 | 2.5 |
| BUV805 | CD4 | 0.2 | 1.25 |
| BV421 | CCR6 (CD196) | 125ul | 4 |
| Pacific Blue | CD49b | 0.5 | 2 |
| BV510 | MHC II | 125ul | 4 |
| BV605 | Ly6C | 125ul | 4 |
| BV711 | CD8a | 125ul | 4 |
| BV785 | F4/80 | 125ul | 4 |
| FITC | CD19 | 0.5 | 0.5 |
| Spark Blue 550 | CD3 | 0.5 | 2 |
| PerCP | CD45 | 0.2 | 1.25 |
| PerCP/Cy5.5 | CD11b | 0.2 | 1.25 |
| PE | CD25 | 0.2 | 4 |
| PE/Dazzle | NK1.1 | 0.2 | 1.25 |
| PE/Cy7 | CCR2 (CD192) | 0.2 | 4 |
| APC | CCR4 (CD194) | 0.2 | 4 |
| AlexaFluor 700 | FOXP3 | 0.5 | 0.12 |
| APC/Fire 750 | Ly-6G | 0.2 | 2.5 |

**Supplemental Figure 1: Gating Strategy** The gating tree was set as follows. *A*: FSC/SSC (represents the distribution of cells in the light scatter based on size and intracellular composition, respectively) to *B*: FSC-H/FSC-A (excludes events that could represent more than 1 cell) to *C*: SSC-H/SSC-A (excludes events that could represent more than 1 cell) to *D*: live gate (live/dead Blue negative, which represents the fraction of viable cells within the sample analyzed) to *E:* SSC-A/CD45 positive (immune cell lineage marker). To identify myeloid immune cells, we gated from E to *F*: FITC-CD19/SparkBlue-CD3 (CD3 – gate for non-T-cell lineage) and J: BV737-CD11c/BV510-MHCII (Dendritic cells). Then F to *G*: PE-Dazz-BVle-NK1.1/PerCP/Cy5.5-CD11b (natural killer cells) and *H*: APCFire700-Ly6G / BV510 – MHCII. Gate in H to *I*: F4/80-BV785 / PerCP/Cy5.5 – CD11b (Macrophage). Then J to K: APCFire700-Ly6G/PerCP/Cy5.5-CD11b (Neutrophils) to BV605-Ly6C/PE/Cy7-CCR2 (Myeloid derived suppressor cells). To identify lymphoid immune cells we gated from E to M: FITC-CD19/SparkBlue-CD3 (B-cells and CD3+ gate for T-cell lineage) to N: BV711-CD8/BUV805-CD4 (CD8+ T-cells and CD4+ T-cells) to O: Alexa700-FoxP3/PE-CD25 (T-regulatory cells) and P: BV421 – CCR6/ APC-CCR4 (TH17).

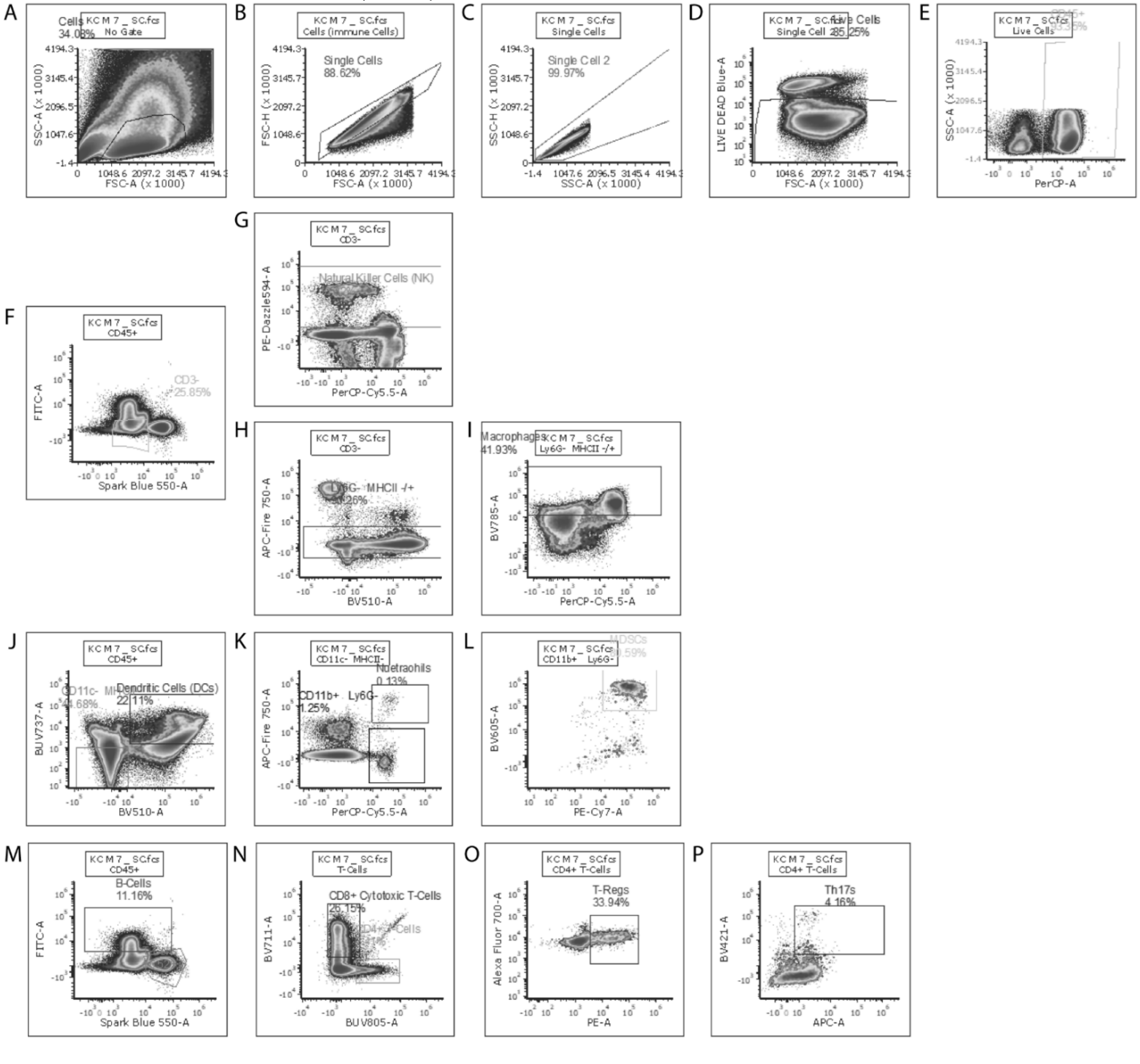

**Supplemental Figure 2: No statistically significant differences in infiltrating dendritic cells, neutrophils and myeloid derived suppressor cells between KC and $A^{-/-}$KC.**

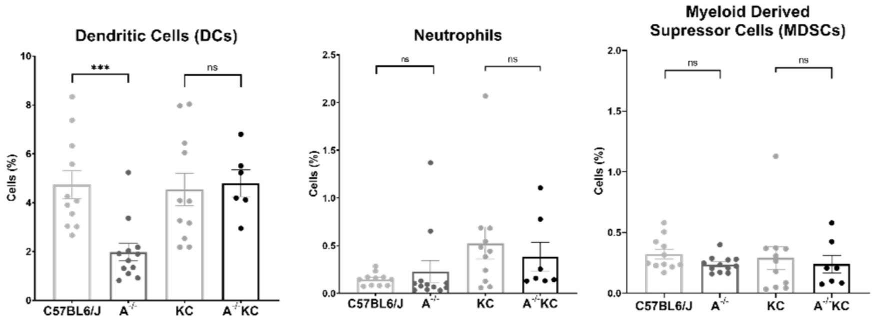

**Supplemental Figure 3: An Increase in immune cell populations and a decrease in fibroblasts found in the A$^{-/-}$KC pancreas through single-cell RNAseq.** A comparative UMAP showing the number of cells in their clusters found in A$^{-/-}$KC pancreas compared to KC pancreas (A). Further confirming these data using a bootstrap comparison method that shows what data would remain significant after 1000 tests, we find that B-cells are most significantly increased in the A$^{-/-}$KC pancreas while fibroblasts are most significantly decreased. The red dots indicate a p-value <0.05 and absLog2FD >0.58 (B).

**A**

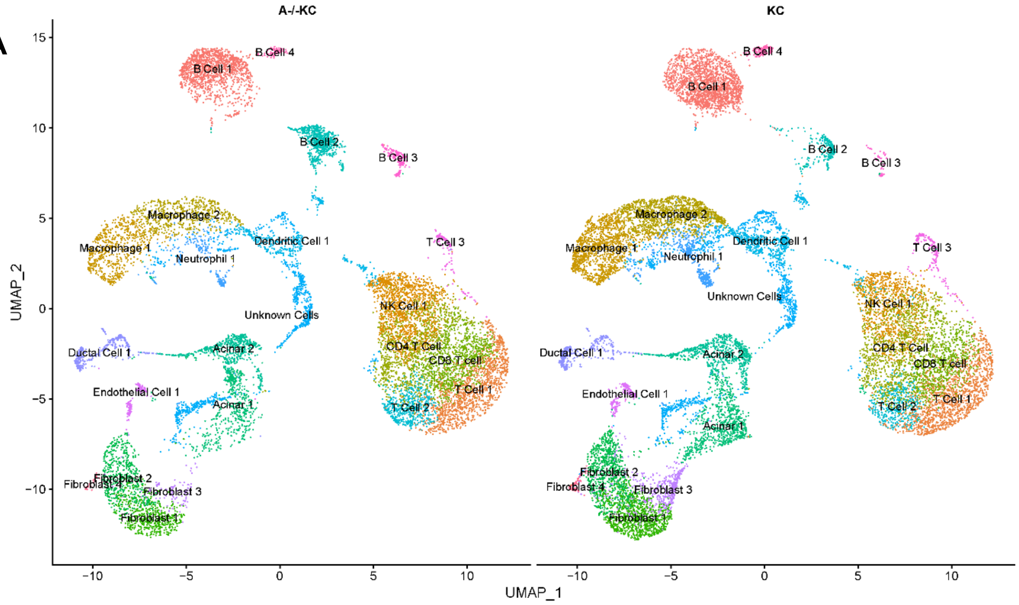


**B**

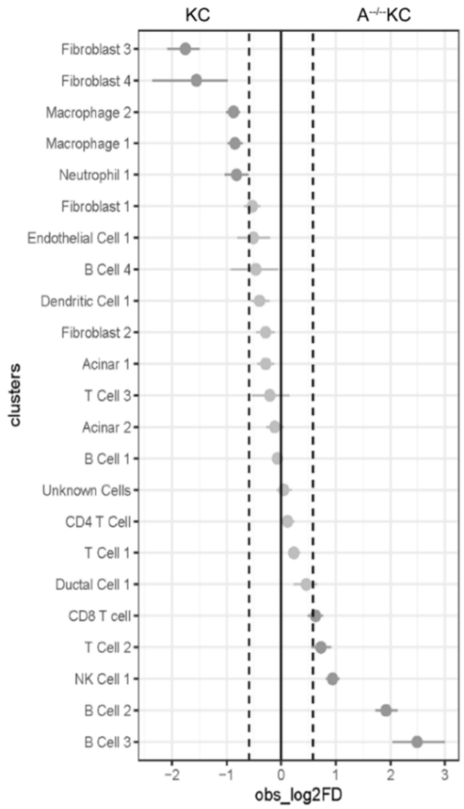
